# Meta-analysis reveals pathway signature of Septic Shock

**DOI:** 10.1101/051706

**Authors:** Samanwoy Mukhopadhyay, Abhaydeep Pandey, Pravat K Thatoi, Bidyut K Das, Balachandran Ravindran, Samsiddhi Bhattacharjee, Saroj K Mohapatra

**Affiliations:** National Institute of Biomedical Genomics, 741251, Kalyani, India; Institute of Life Sciences, NALCO Square, 751023,Bhubaneswar, India; Sriram Chandra Bhanja Medical College and Hospital, 753007,Cuttack, India

## Abstract

Septic shock is a major medical problem with high morbidity and mortality and incompletely understood biology. Availability of genome-wide expression data from different studies on septic shock empowers the quest for hitherto unidentified pathways by integration and meta-analysis of multiple data sets. Electronic search was performed on medical literature and gene expression databases. Selection of studies was based on the organism (human subjects), tissue of origin (circulating leukocytes) and the platform technology (gene expression microarray). Gene-level meta-analysis was conducted on the six selected studies to identify the genes consistently differentially expressed in septic shock. These genes were then subjected to pathway analysis. The identified up-regulated pathway *hsa*04380 (Osteoclast Differentiation) was validated in an independent cohort of patients. A simplified model was generated showing the major gene-modules dysregulated in SS.

## Introduction

Septic shock (SS) is a serious medical condition that claims many lives every year worldwide. Approximately 2% of the patients admitted to the hospital are diagnosed with SS. Of these patients, half are treated in the intensive care unit (ICU), representing 10% of all ICU admissions (*2, 29*). Approximately 40-60% of the SS patients die within 30 days (*2*). The number of cases in the USA exceeds 750,000 per year (*2*), but the incidence of SS is largely unknown in those parts of the world with scarce ICU care. Extrapolating from treated incidence rates in the USA, Adhikari et al. estimated up to 19 million cases worldwide per year (*1*). However, the true incidence is expected to be far higher. Incomplete grasp of SS biology is compounded by the lack of specific drug for treating the condition. With a number of failed clinical trials, there is urgent need for new directions in research (*5*). Genome-wide expression profiling offers a complete picture of the condition and enables identification of genes and pathways of diagnostic, prognostic or therapeutic relevance (*37*).

The purpose of this study was to investigate genome-wide host response to SS by combining the power of multiple studies. Analysis was performed at two levels: genes and gene sets. A “gene set” or pathway consists of a set of functionally related genes, and provides higher-order information about gene expression and valuable insights into the biology of a disease. Accordingly, we have laid emphasis on discovery of pathway(s) enriched among the genes consistently differentially expressed among the multiple data sets from studies on SS.

A biological process involves a group of genes. The principle of enrichment analysis is that if a biological process is abnormal in a given condition, the co-functioning genes should have a higher (enriched) potential to be implicated as a relevant group. Because the analytic conclusion is based on a group of relevant and functionally related genes instead of on individual genes, it increases the likelihood of identifying the biological process(es) pertinent to the disease condition under study. A variety of methods are available for testing differential expression of gene sets. We started with the most popular method that takes a list of differentially expressed genes and tests whether the gene set is over-represented in this list. This was followed up and validated by other methods to discover a pathway that is consistently perturbed in SS.

## Methods

### Search strategy and selection criteria

We searched the popular online database PubMed with the search string (“Systemic Inflammatory Response Syndrome”[MeSH] OR “septic shock”[MeSH] OR “Shock, Septic”[MeSH] OR “Endotoxemia”[MeSH]) AND (“gene expression profiling”[MeSH] OR “transcriptome”[MeSH] OR “microarray analysis”[MeSH] OR “Oligonucleotide Array Sequence Analysis”[MeSH]) with “Human” filter. Additionally, we searched the following gene expression databases: (1) National Centre for Biotechnology Information Gene Expression Omnibus (GEO) and (2) European Bioinformatics Institute ArrayExpress. All the queries were made on the 30th Aug, 2013. Entries from the gene expression databases were then cross-referenced with publications retrieved from PubMed. Selection of studies was based on the organism (human subjects), tissue of origin (circulating leukocytes) and the platform technology (gene expression microarray). The process of selection of the six studies of SS is described in Fig. 1. Only data sets published as full reports and meeting the previous three criteria were selected (Table 1). These data sets were subjected to systematic analysis, as described below.

**Table 1:** SS Study Characteristics. The table shows characteristics (such as sample size, study design, clinical parameters for inclusion criteria and details about the platform technology used to generate the data) of the six selected gene expression studies of septic shock

| Study ID | Author-Year | PMID | Study Design | Total No. of Samples | Mean Age (yr) | Clinical Setting | Inclusion Criteria | Control Group | Tissue | Sample collection and RNA isolation method | Platform | Data normalization |
| --- | --- | --- | --- | --- | --- | --- | --- | --- | --- | --- | --- | --- |
| GSE4607 | Wong2007<br>Cvijanovich2008<br>Wong2012 | 17374846<br>18460642<br>23107287 | Observational study | 84 | Children <10 years of age | PICU | Septic Shock | Healthy controls | Whole blood | PAXgene | Affymetrix Human Genome U133 Plus 2.0 Array | RMA |
| GSE8121 | Shanley2007 | 17932561 | Longitudinal | 75 | Children <10 years of age | PICU | Septic Shock | Healthy controls | Whole blood | PAXgene | Affymetrix Human Genome U133 Plus 2.0 Array | RMA |
| GSE9692 | Cvijanovich2008 | 18460642 | Retrospective observational study | 45 | Children <10 years of age | PICU | Septic Shock | Healthy controls | Whole blood | PAXgene | Affymetrix Human Genome U133 Plus 2.0 Array | RMA |
| GSE13904 | Wong2009 | 19325468 | Prospective observational study | 124 | Children <10 years of age | Paediatric wards | Septic Shock | Healthy controls | Whole blood | PAXgene | Affymetrix Human Genome U133 Plus 2.0 Array | RMA |
| GSE26378 | Wynn2011 | 21738952 | Prospective observational study | 103 | Children <10 years of age | Paediatric wards | Septic Shock | Healthy controls | Whole blood | PAXgene | Affymetrix Human Genome U133 Plus 2.0 Array | RMA |
| GSE26440 | Wong2009 | 19624809 | Prospective observational study | 130 | Children <10 years of age | Paediatric wards | Septic Shock | Healthy controls | Whole blood | PAXgene | Affymetrix Human Genome U133 Plus 2.0 Array | RMA |

**Figure 1:**
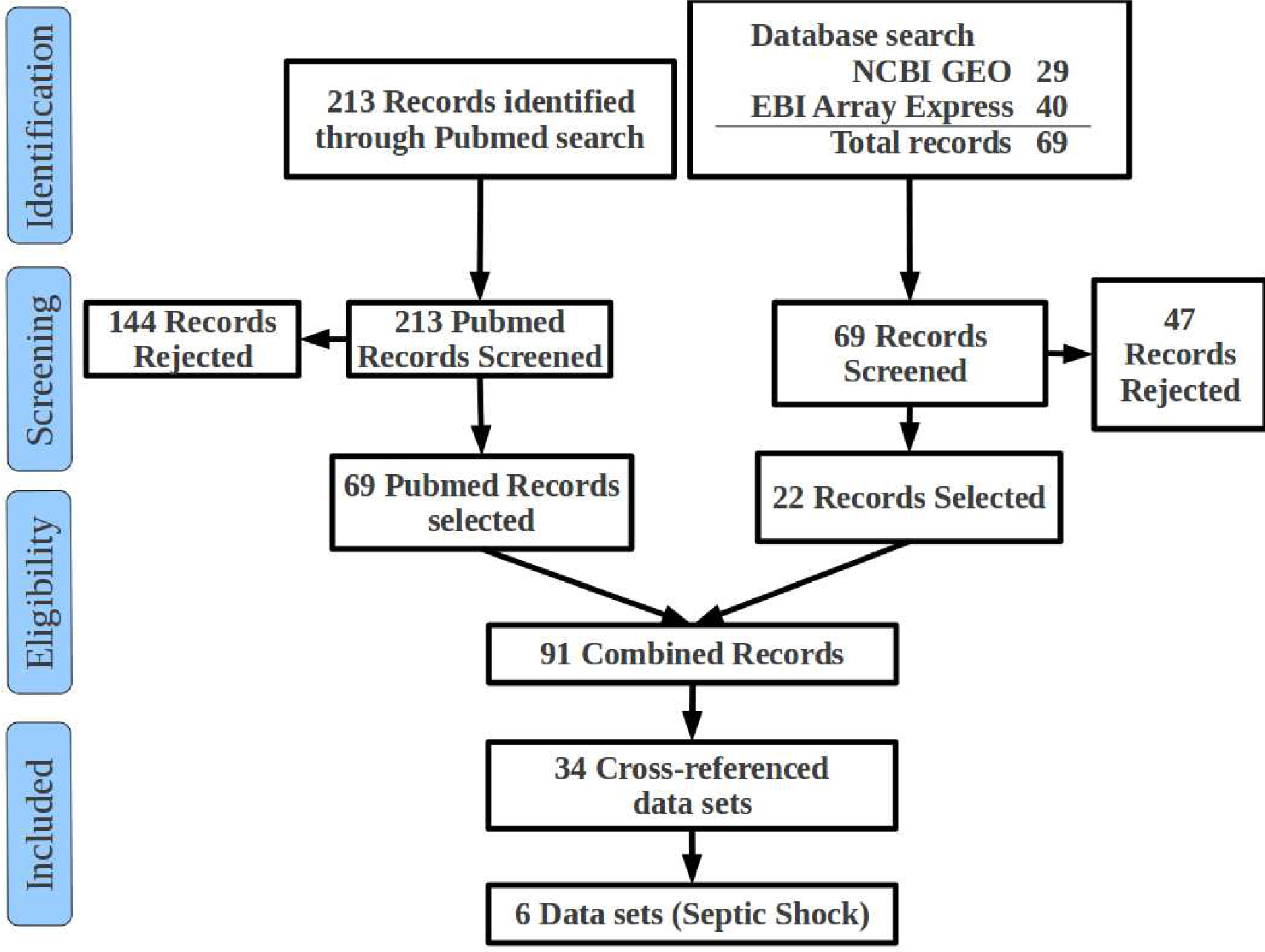
Selection of studies. Electronic search on the Gene expression and Literature databases was performed to identify the 213 Pubmed records and 69 gene expression data sets. These records were manually screened to select the studies that were done in human subjects, on circulating leukocytes and gene expression profiling done using gene expression microarray platform, resulting in 91 combined (Pubmed and expression data) records. Duplicate records were removed, and records without data from human sepsis or SS, were deleted. One record was added from a source outside those mentioned above. Of the resulting 34 data sets (cross-referenced with Pubmed articles), six were derived from studies on SS. These six data sets were selected for meta-analysis.

### Pre-processing of data

Series matrix files (containing normalized gene expression data) for all six studies were downloaded and read with the function *getGEO* of the bioconductor package “GEOquery” (*7*). All data were transformed to logarithmic scale (base 2). The function *nsFilter* of “genefilter” package (*8*) was applied to remove the probe sets with incomplete or duplicate gene annotation, and those carrying low expression signal. Each probe set ID was mapped to the corresponding Entrez gene ID. Genes common to all six studies were included in the analysis. These common genes were extracted and all further analyses were performed only on this list of common genes across all the studies. Some of the studies included samples from cases other than SS in the data set. However, these samples were excluded from analysis. For each study, control samples were retained for comparison with the cases of SS.

### Identification of differentially expressed genes

For all genes within each study, we applied a Welch’s 2-sample t-test (*36*) to compare expression values in samples of SS with that in samples of control subjects. A single overall p-value was then computed for each gene by meta-analyzing individual study level t-statistic using the following procedure.

We first transformed each individual t-statistic to a Z-score Z_*i*_ retaining its sign by using the quantile transformation:

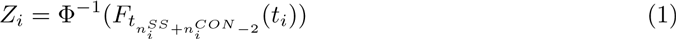

We then followed a fixed effects meta-analysis (*11*) approach, where the Z-scores are combined using an optimal linear combination with weights equal to square root of the effective sample size of each study (i.e. the harmonic mean of case and control sample sizes). This gave the combined meta-analyzed Z-score Z_*meta*_.

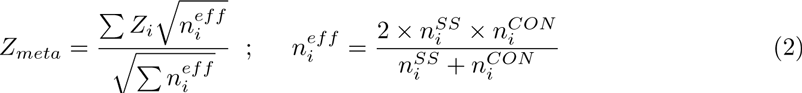

Thus the final overall 2-sided p-value *p*_*meta*_ was obtained by:

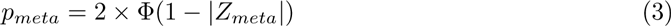

The p-values for all genes were then adjusted for multiple testing using the method of Benjamini and Hochberg (1995) (*3*) by a call to the function *p. adjust* of the R package “stats” (*28*). For each gene, mean log-fold change in gene expression was computed by averaging the log-fold changes over the six studies. Differential expression was detected by applying two thresholds: p-value < 0.01 and 2 fold or greater change in expression in SS compared to control.

### Over-Representation Analysis (ORA)

The set of up-regulated genes was then subjected to over-representation analysis by applying hypergeometric test which is briefly described here. Let us consider two lists of genes: the first list being the set of up-regulated genes, and the second, genes that are part of a given KEGG pathway (*14*). The task is to find out if genes belonging to this pathway are also likely to be part of the list of up-regulated genes. This is captured in a 2×2 contingency table as shown in Table 2.

**Table 2:**
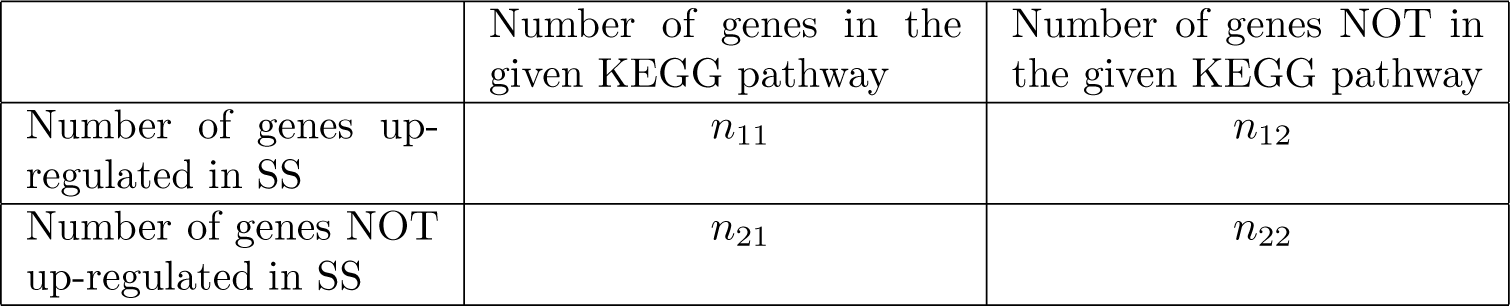
A 2 × 2 contingency table. The table shows the four quantities of interest while estimating if a pathway is over-represented (enriched) among the set of differentially expressed genes.

|  | Number of genes in the<br>given KEGG pathway | Number of genes NOT in<br>the given KEGG pathway |
| --- | --- | --- |
| Number of genes up-regulated in SS | $n_{11}$ | $n_{12}$ |
| Number of genes NOT<br>up-regulated in SS | $n_{21}$ | $n_{22}$ |

Over-representation analysis tries to capture whether n_11_ and n_22_ are larger than expected relative to the other two cell numbers; in other words do genes belonging to the KEGG pathway also “tend to be in the up-regulated category.”

Hypergeometric testing was applied to detect if a significant association exists between the two attributes of a gene: being up-regulated in SS and being member of a particular pathway. A significant association was detected at threshold of p < 0.001. Over-representation analysis was implemented by applying the function *hyperg* of the “Category” package (*9*).

### Gene Set Enrichment Analysis (GSEA)

GSEA focuses on cumulative changes in the expression of multiple genes as a group, which shifts the focus from individual genes to groups of genes. By looking at several genes at once, GSEA identifies pathways with coordinated change in expression of the member genes. The following steps were taken. First, genes were mapped to pathways on an incidence matrix. Then *gseattperm* function from the “Category” (*9*) package was used for the permutation test. The function received the following as input: the gene expression data, sample phenotypic data, and the incidence matrix representing the gene sets of interest. Then it created a version of the data set with phenotype labels randomly scrambled, producing the corresponding ranked list, and computed the enrichment score (ES) of the gene set for this permuted data set. This process of scrambling the labels and re-computation of ES was repeated 10000 times to produce an empirical null distribution of the ES scores. The observed ES for each pathway was compared against the null distribution. Pathways with observed ES very different (p = 0.0) from the null distribution were considered significant. For each pathway a meta-analyzed GSEA p-value was obtained using Fisher’s p-value product method (*38*).

### Signaling Pathway Impact Analysis (SPIA)

Impact Factor analysis combines both over-representation analysis and functional class score (gene set enrichment analysis) approach, taking into account fold-change in gene expression, gene-gene interactions such as activation and inhibition, and the topology of the pathway (*33*). By combining two different evidences (one from the analysis using the hypergeometric model and the other from the probability of perturbation that takes the pathway topology into account), a pathway-level score was computed. For each gene, t-test between SS and control was performed. Log-fold changes of the differentially expressed (p < 0.05) genes were passed to the function *spia* of the package “SPIA” (*33*). This was repeated for each study separately, each time returning a list of pathways and associated p-values. For each pathway, Fisher product of the six p-values (one for each study) was calculated to return one p-value for pathway. The ten most significantly perturbed pathways were considered.

### Validation cohort

Blood samples were collected from the patients of SS and healthy subjects after obtaining approval from the Institutional Ethical Committees of the National Institute of Biomedical Genomics, Kalyani, India and SCB Medical College Hospital, Cuttack, India. All the methods were carried out in accordance with the approved guidelines. Informed consent was obtained from all subjects who participated in the study. 12 healthy control and 9 cases of SS were included in the study. Whole blood was collected from the subjects in PAXgene Blood RNA tubes (BD/Preanalytics) and incubated at ambient temperature for at least 2 hours. RNA was isolated using PAXgene Blood RNA kit (Qiagen) as per manufacturer’s instructions. Total RNA yield and quality were checked in Nanodrop 2000 spectrophotometer and Agilent Bioanalyzer RNA Nano 6000 chip. RNA samples with a RIN number ≥ 6 were converted to cRNA and hybridised onto either HumanHT-12 v4 BeadChip (Illumina) or Human Gene 2.0 ST Arrays (Affymetrix). Gene expression intensities were read using the standard protocol provided by the manufacturer.

Raw data files from the platform HumanHT-12 v4 BeadChip (Illumina) were read with the function *read.ilmn* of the package “limma” (*32*). After reading the raw data the background correction and normalization using the control probes were done by the help of function *neqc* of the package “limma”. For the samples processed on Human Gene 2.0 ST Arrays (Affymetrix), RMA-normalized data were read with the function *read.table*. Each probe ID was mapped to its corresponding Entrez gene ID from the gene symbol by the help of “org.Hs.eg.db” package. In case of duplicated/multiple probes for a single Entrez gene id, the probe with the highest variance was retained (and others dropped from further analysis using the function *nsFilter* of the package “genefilter”). Correction of batch effect was performed with the *ComBat* function of “sva” (*41*) package. Genes of the pathway hsa04380 were extracted and mean gene expression computed. Scatter plot of SS versus control was generated for visual inspection of the data.

### Permutation test for enrichment

We performed a permutation-based enrichment test to provide evidence for overall up-regulation of the pathway hsa04380 in SS (validation cohort). For this, we used the function *permutationTest* of the package“resample” (*39*) to calculate the permutation-based p-value accounting for correlation among the pathway genes. First, we calculated the proportion of significantly up-regulated (p < 0.05) genes in the pathway by using a two-sample t-test to test for up-regulation of each pathway gene. This was observed to be 0.447. Next, we reshuffled the sample groups (i.e. case control status) 100000 times and similarly calculated the proportion of up-regulated genes for each permutation replicate. Finally, the permutation-based p-value was obtained as the proportion of replicates where the simulated proportion was greater than the observed value.

Gene-level analysis: We then looked at the individual genes in the pathway that were significantly up-regulated after multiple-testing correction at an FDR level of 0.05. We further filtered the resulting gene list to include only genes that showed a high fold-change (2 or more) and were expressed in significant amounts (intensity of 100 or more).

### Software used

All analyses except Ingenuity Pathway Analysis (described later) were done with R version 3.1.0 implemented under GNU/Linux operating system on a 32-bit i686-computer system. The analysis work flow is presented in Fig. 2.

**Figure 2:**
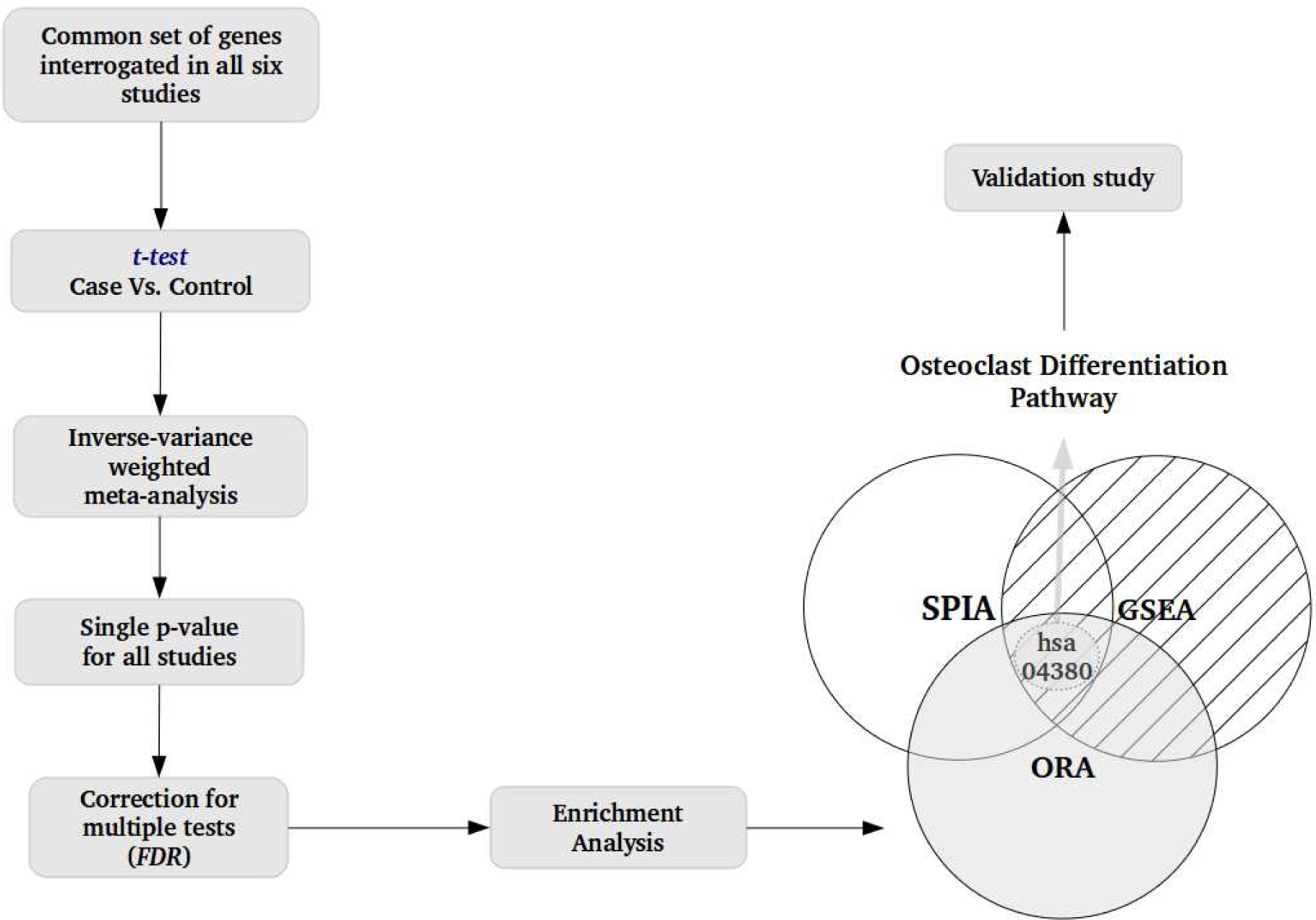
Overview of the analysis work-flow. First, gene-level analysis was performed by comparing SS with control. The p-values from multiple studies were combined to compute a single p-value per genes. The p-values were adjusted for multiple testing (FDR). 200 genes with low p-value (p < 0.001) and large fold change (2 or greater) were considered significantly differentially expressed (up-regulated) in SS. The 200 up-regulated genes were then subjected to three pathway analysis methods [Over-Representation Analysis (ORA), Gene Set Enrichment Analysis (GSEA) and Signaling Pathway Impact Analysis (SPIA)]. A single pathway (hsa04380 - Osteoclast Differentiation) was returned as the common result of the three approaches. Up-regulated gene expression of hsa04380 in SS was validated in an independent patient cohort.

### Ingenuity Pathway Analysis of the up-regulated genes

Mean log-fold change in expression was calculated for the 200 genes up-regulated in SS, and submitted to the Ingenuity Pathway Analysis (IPA®, QIAGEN Redwood City, www.qiagen.com/ingenuity). Core IPA analysis was performed without applying any threshold or cutoff. IPA performed a right-tailed Fisher’s Exact Test and calculated a p-value for over-representation of *a priori* defined biological pathways in the data submitted. It also corrected the p-value to account for multiple testing by Benjamini-Hochberg method (FDR). IPA generated molecular networks with algorithmically generated pathways which were over-represented in the submitted data. Similarly it identified diseases and biological functions that were over-represented in the data. The three broad categories returned by IPA were interpreted.

### Availability of data and materials

The dataset(s) and the R code supporting the conclusions of this article are available under the project nibmgss in the SourceForge repository https://sourceforge.net/projects/nibmgss.

## Results

### Selection of studies and meta-analysis

By systematic search of literature, NCBI GEO and other gene expression databases, we selected six studies of SS (Fig. 1, Table 1). Normalized gene expression data of the studies were retrieved from GEO and analyzed to detect differentially expressed genes (Fig. 2). Differential expression was measured in terms of both log-fold change and p-value. For each gene, SS was compared with control and the six p-values were combined and adjusted for multiple testing (detailed meta-analysis procedure provided in the Methods section) to generate a single p-value per gene. For each gene, the six log-fold changes were averaged to produce a single log-fold change. Using stringent criteria (adjusted p-value < 0.01, fold change of 2 or more), we discovered 200 genes that were consistently up-regulated in SS. We did not discover any gene that was down-regulated at this level of significance, i.e., p < 0.01 and fold change of half or less (Supplementary Fig. S1).

### The 200 genes up-regulated in SS

These included genes associated with inflammation and innate immunity (SERPINB1, S100A12, SLC11A1, TLR5, PADI4, CLEC5A, TLR8, LILRA5), critical illness (ANKRD22, CEACAM1, CLEC5A, G0S2, BCL-6, MMP8, TREM1, CCL4), anti-microbial activity (ADM, BP1, DEFA4, RNASE3), complement and coagulation systems (C1QB, F5, CR1, C3AR1, CR1, C3AR1, CD59, SERPINA1), carbohydrate and lipid metabolism (PFKB2, PFKB3, SLC2A3, GYG1, PYGL, ACSL1, HGF, FAR2, DGAT2), and others. While many of these systems are known to be involved in SS, there were some surprises, such as, up-regulation of genes of bone metabolism (CPD, MAPK14, FOSL2, IL1R1, MAP2K6, SOCS3, OSCAR, SIRPA, ALPL, BST1, CA4, TNFAIP6) in SS (Supplementary Table S1). With strong evidence in favour of altered expression at gene level, we proceeded to analyse the data at pathway (gene set) level.

### Pathway analysis: ORA, GSEA, SPIA, IPA

A pathway is a set of genes that participate in a single functional module. When a large number of genes in the same pathway are perturbed (or up-regulated, in this case), then the pathway itself may be inferred to be perturbed. Secondly, small changes in gene expression are usually not detectable with standard methods. However, if the changes happen in a co-ordinated fashion over a set of functionally related genes (i.e., a pathway), then the cumulative effect of the small gene-level changes shall result in a large and detectable perturbation at the pathway-level. Lastly, the gene set in a pathway has a specific network structure or topology, such that, the nodes proximal to the point of origin (for example, the signal from the membrane), has a greater regulatory impact compared to the distal nodes. All these considerations were taken into account in the current analysis.

Kyoto Encyclopedia of Genes and Genomes (KEGG) (*14*) is a database of gene sets (pathways) arranged by function. Each pathway consists of multiple genes and a gene may belong to more than one pathway. We reasoned that, for a pathway to be up-regulated in SS, a large number of genes in the pathway need to be up-regulated. In other words, we sought to find the pathway(s) that is (are) significantly “enriched” with the genes in our list of 200. Using the method of “Over-Representation Analysis” (ORA; described in detail in the Methods section), three pathways were identified: hsa04610 - Complement and Coagulation Cascades, hsa04380 - Osteoclast Differentiation, and hsa05202 -Transcriptional Misregulation in Cancer (Supplementary Table S2). The complement system is a proteolytic cascade in blood plasma participating in innate immunity. When activated, it leads to recruitment of inflammatory cells and elimination of pathogens. Dysregulation of the complement system (especially activated C3 and C5) is known to occur in SS (*23*). Similarly, blood coagulation is an essential defense mechanism for blocking off the pathogen and preventing spread of inflammation, in the initial phase. However, the inflammatory state found in SS alters the hemostatic balance in favour of a procoagulant state, with disseminated intravascular coagulation (DIC) (*17*). The other two pathways are not commonly associated with sepsis or SS, although osteoclasts arise from differentiation of blood monocytes and cause bone resorption, which is markedly increased in critically ill patients (*31*). The third pathway hsa05202 consists of many genes that are transcriptionally altered in different malignancies.

We next applied two other tools, “Gene Set Enrichment Analysis” (GSEA) and “Signaling Pathway Impact Analysis” (SPIA), that are methodologically independent of ORA, and provide additional support to our finding (Fig. 2). GSEA calls upon a global (i.e., genome-wide, not limited to any pre-selected list) search strategy to detect the KEGG pathway (s) with significant up-regulation in SS compared to control. SPIA combines elements of ORA and GSEA, with attention to gene-gene interactions and pathway topology. There was a single pathway (hsa04380) that was common to the results from the three approaches (ORA, GSEA, SPIA). Further, we performed Ingenuity Pathway Analysis (IPA) with the top 200 up-regulated genes, that returned a list of pathways. Two pathways among the top ten were related to bone loss in Rheumatoid Arthritis. This and the presence of other pathways, such as, p38 MAPK signaling, complement system and TLR signaling, were similar to the results from KEGG analysis. IPA discovered the following diseases and functions enriched among the up-regulated genes: Infectious Disease, Connective Tissue Disorder, Inflammatory Diseases and Skeletal and Muscular Disorders. The network with significant biological function termed “infectious disease, inflammatory response, connective tissue disorders” was discovered to be differentially regulated in SS (Supplementary Fig. S2). Over all, IPA results were similar to the pathway analysis with KEGG, and generally consistent with the finding of altered skeletal function in SS. Thus, multiple analytic approaches, ORA, GSEA, SPIA and IPA, together revealed hsa04380 (Osteoclast Differentiation) with unequivocal transcriptional up-regulation in SS. We then sought to validate the pathway up-regulation in an independent set of SS cases.

### Validation of hsa04380 in an independent cohort

Batch-corrected whole blood gene expression data (Supplementary Fig. S3) of the pathway hsa04380 was obtained from an independent set of human subjects (Supplementary Table S3). Mean expression values (for the two groups: control and SS) were calculated and a scatter-plot generated (Fig. 3). In this plot, each point corresponds to a single gene. The points near the identity line (the diagonal in the Figure), correspond to genes with similar expression level in control and SS groups. The genes that are up-regulated in SS are expected to be significantly deviated from the diagonal toward the SS axis. Indeed, for most of the genes, there is much higher expression in SS, as shown in Fig. 3. Additional evidence for up-regulation of the pathway hsa04380 was obtained from the permutation-based enrichment test, where we resampled (reshuffled) the case/control labels of the samples and calculated a permutation-based p value for the proportion of up-regulated genes in the pathway. We observed that up-regulation of the pathway was highly significant (permutation p = 0.00028; 100000 replicates). On looking at individual genes, we found that 60 genes were FDR significant and 25 genes passed the expression filters. Supplementary Fig. S4 shows the boxplots of these highly significant genes in the validation cohort.

**Figure 3:**
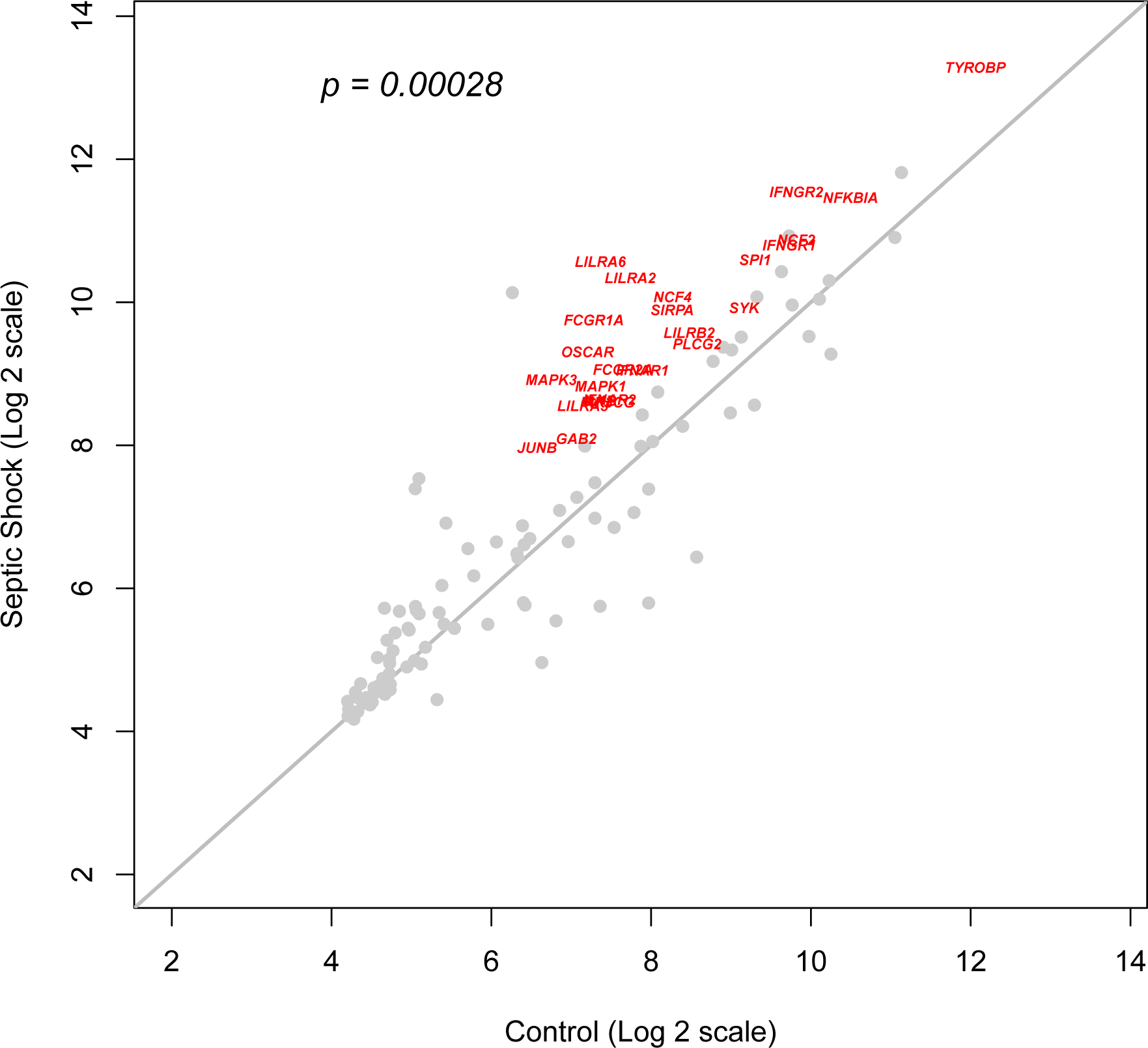
Up-regulation of Osteoclast Differentiation pathway in septic shock. Scatter plot of hsa04380 (Osteoclast Differentiation) pathway genes from expression data generated in an independent validation cohort. Each point corresponds to a single gene, and its coordinates reflect mean gene expression in healthy controls (x-axis) and SS (y-axis). The points close to the identity line (the diagonal in the figure), correspond to genes with similar expression level in the control and the SS groups. Points away from the diagonal represent genes that are differentially expressed in SS (up-regulated if the points are in the upper half of the plot area; down-regulated otherwise). Many of the genes are up-regulated in SS as shown by a large number of points deviated from the diagonal toward the SS-axis. The up-regulation of these genes in SS is statistically significant (p = 0.00028, permutation test for enrichment). Additional testing was performed for each gene (unpaired t-test between control and SS) leading to selection of individual genes in the pathway that were significantly up-regulated after multiple-testing correction at an FDR level of 0.05. An expression filter was applied to identify the genes that showed a high fold-change (2 or more) and were expressed in significant amounts (intensity of 100 or more). These genes have been shown in red on the plot.

The 25 highly significant genes of the pathway are well-connected in the pathway map, and known to be expressed in neutrophils and monocytes, which are the two abundant sub-types of leukocytes. These 25 genes are not only sgnificantly up-regulated in the validation cohort but also significantly (p< 0.05; in gene level metaanalysis) up-regulated in the Discovery cohort (6 SS datasets).

### Significant genes of the pathway in SS

Of these 25 key genes of this pathway (Supplementary Table S4), there were genes whose encoded proteins form a signal transduction cascade from the plasma membrane to the nucleus. There were genes coding for proteins situated on the plasma membrane (IL1R, TNF*α*, TGF*β*), or belonging to MAP Kinase group (Rac1, MKK6 and p38), AP1 transcription factor complexes (JUNB, FOSL2). This suggested that a signal starting from the membrane may be transduced through the MAP kinase subnetwork to the nucleus culminating in transcriptional regulation of osteoclast differentiation-specific genes. The list also included other genes (OSCAR, TREM2, SIRPB1, NFATC1) that suggested a parallel path from the membrane leading to up-regulation of the transcription factor PU.1 and expression of genes for osteoclast differentiation. Based on these 25 genes, a simple model of osteoclast differentiation in SS was proposed (Fig. 4).

**Figure 4:**
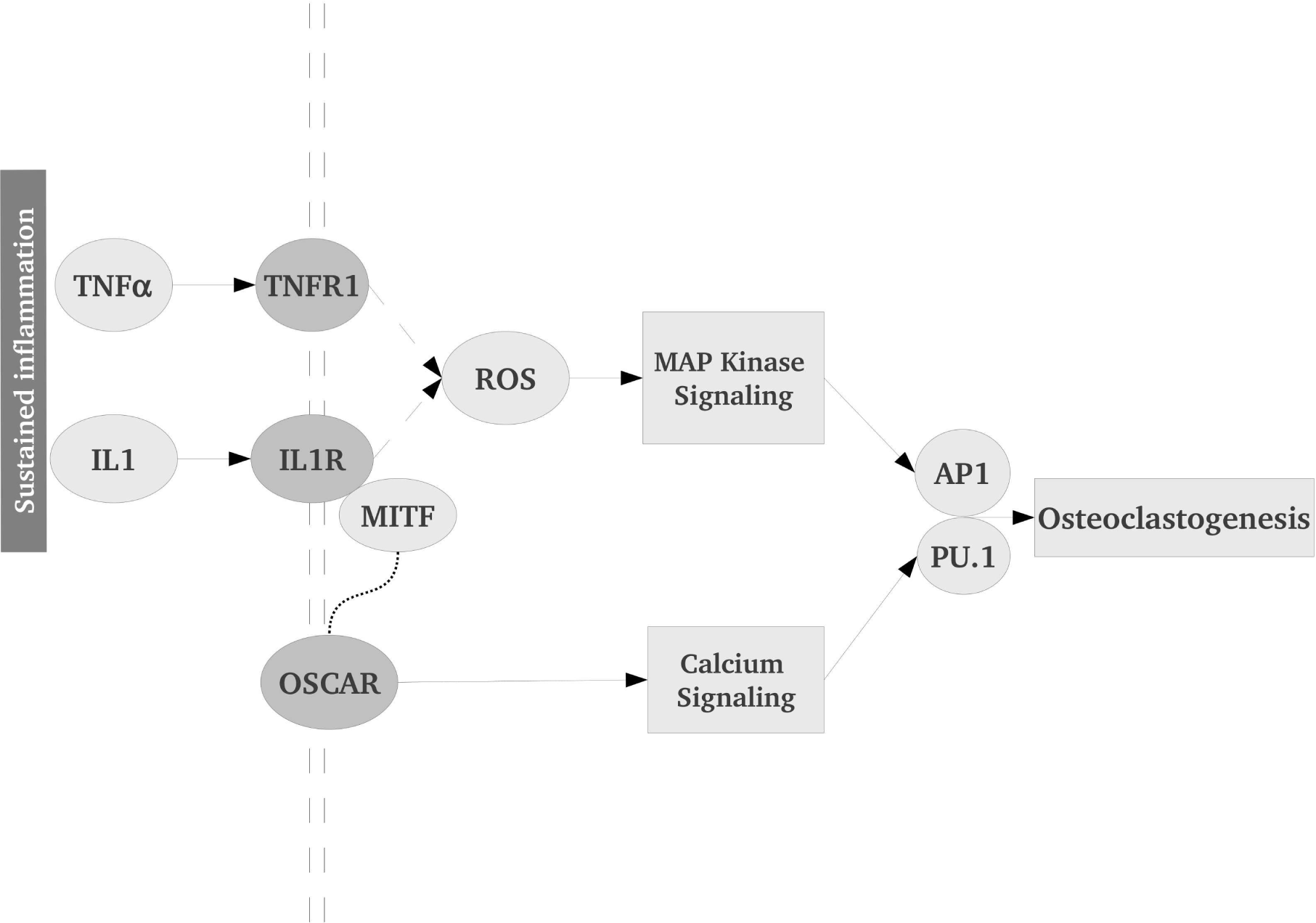
A simplified model of the pathway. A simplified model of the Osteoclast Differentiation Pathway up-regulation in SS. Sustained inflammation leads to activation of osteoclast differentiation through proinflammatory cytokines TNFκand IL1, followed by MAP Kinase and Calcium signaling. Reactive Oxygen Species (ROS) play a major role in activation of MAP Kinase. OSCAR is an important receptor on the membrane that causes signal transduction via calcium-signaling. AP1 and PU.1, two important transcription factors, integrate multiple signals and induce expression of osteoclastogenesisspecific genes.

Since our analysis was performed on whole blood expression data, we were also interested in the cellular and plasma distribution of the gene products. A literature search for tissue specific expression revealed that many of these genes are expressed in neutrophils and monocytes, two important leukocyte sub-types. Additionally, almost all of the gene products have been reported to be detectable in plasma (Supplementary Table S4).

## Discussion

In contrast with gene-level analysis, pathway analysis strives to identify a perturbed pathway (a set of functionally related genes) whose transcriptional alteration is significantly associated with the given condition. There exist several approaches to this end, such as ORA, GSEA, or SPIA, each with its own advantages (*15*). ORA considers the number of genes differentially expressed, but ignores any value associated (such as fold-change). GSEA considers all genes in a pathway and is more likely to detect coordinated changes in gene expression. SPIA, in addition, takes the topology of the pathway into account. Here, combination of three different methods gives leverage in finding the pathway that is unequivocally altered in SS.

In order to discover the stable blood transcriptional changes in SS, we conducted three different analyses (ORA, GSEA, SPIA) and intersected the resulting lists to obtain a single KEGG pathway (hsa04380 - Osteoclast Differentiation Pathway) as a stable expression signature of SS in circulating leukocytes. We confirmed this result in an independent validation cohort and demonstrated that up-regulation of the pathway is distinct in SS compared to controls. Further, we subjected the list of up-regulated genes to Ingenuity Pathway Analysis and observed broad agreement with this finding. To our knowledge, this is the first time that osteoclast differentiation pathway has been shown to be significantly perturbed in SS.

The skeletal and the immune systems are known to interact with each other. For example, in immune disorders, such as rheumatoid arthritis, the ratio of bone resorption to bone formation is skewed toward the former. This ratio is balanced by the activity of two groups of cells: osteoclasts and osteoblasts. Osteoclasts are specialised bone cells of monocyte-macrophage lineage (*10, 34*) responsible for bone resorption whereas osteoblasts are responsible for bone formation. It was earlier shown that there is increased bone resorption in critically ill patients (*31*). However, our finding suggests that the role of this pathway may be more specific in SS.

We propose a simplified model of transcriptional changes in hsa04380 during SS, as described below and shown in Fig. 4. The Osteoclast differentiation pathway may be conceptualised in terms of interactions among a few key processes or subnetworks. Firstly, pro-inflammatory cytokine-receptor complexes (such as, TNF*α*-TNFR and IL1-IL1R) can cause osteoclast differentiation with the help of MITF and MAPK, NF*κ*B signaling (*16, 18*). In parallel, IL-1 and TNF*α* activate NF*κ*B to up-regulate expression of osteoclast specific genes CD40LG (also known as TRAP), CTSK (cathepsin K) and OSCAR (osteoclast associated receptor), three important regulators of bone mass. It is worth mentioning that IL1-mediated osteoclast differentiation is distinct from the well-known RANKL-RANK signaling mediated activation (*16*). We observed significant up-regulation of CD40LG, cathepsin K and OSCAR, but not RANKL or RANK. The association of TNFα, IL-1, IL-6 and IL-7 has been detected in a variety of chronic inflammatory bone diseases such as rheumatoid arthritis, osteoarthritis, and periodontal diseases. These cytokines are produced by macrophages, lymphocytes, osteoblasts and bone marrow stromal cells under the regulation of NF*κ*B, and also stimulate NF*κ*B signaling in the target cells by positive feedback (*20*). These genes are significantly up-regulated in SS.

The second subnetwork starts with a set of closely related and functionally similar immunoglobulin-like receptors among which the most prominent one is OSCAR (*6*). Each receptor molecule bears ITAM transmembrane domains that transduce signal from plasma membrane to the nucleus through second messenger molecules, such as Syk (a tyrosine kinase) and BLNK, which carry the signal to PLC_γ_, resulting in intracellular calcium release, that is known to activate transcription factors. Two transcription factors PU.1 (a well-known biomarker for osteoclast differentiation) (*30*) and AP1 (JUN, FOSL) (*35*) form a complex that results in transcription of osteoclast-specific genes in the precursor cells. Both PU.1 and AP1 are significantly up-regulated in SS.

It is also interesting, that AP1 is involved in cellular proliferation, transformation and death. There is also a report that apoptotic cell death (mostly of lymphocytes) is observed in patients suffering from SS (*42*). An alternative interpritation would suggest prognosis towards severe diease phenotype (i.e., SS) by AP1-induced excessive lymphocyte apoptosis.

Thirdly, the MAPK module (includes genes such as MAP2K6, JNK, p38 along with Reactive Oxygen Species (ROS) genes NCF2, NCF4) plays an important role in activating AP1 transcription factor complex and regulates genes responsible for the differentiation process (*22*). Generation of reactive oxygen species and MAP kinase pathway signaling play a critical role in this pathway. IL1R1, a prominent mediator involved in many cytokine-induced immune and inflammatory responses, and genes of MAP kinase pathway (MAPK14, MAPK26, FOSL2, IL1R1, CA4), all associated with inflammation, are up-regulated in SS. The signalling molecule p38, known to be essential for osteoclast differentiation (*24*), is sensitive to activation by ROS in bone marrow monocyte-macrophage lineage (BMM) cells (*21*). Mitochondrial dysfunction in SS causes generation of various reactive oxygen (and nitrogen) species, contributing to activation of MAP kinase pathway and subsequent osteoclastogenesis (*21*). Although little attention has been paid to the role of ROS in differentiation of macrophages and monocytes into osteoclasts, we observed that MAP kinase pathway and genes associated with ROS generation are significantly up-regulated in SS. Additionally, both MKK6 and p38 (up-regulated in SS) cause transdifferentiation of neutrophils to monocyte or dendritic cells, which is of relevance during inflammation (*13, 19*).

In most studies, discharge from the ICU is considered the survival end-point. However, the survivors themselves are prone to many long term complications including musculoskeletal ones. By rigorous analysis, we show that patients of SS have increased osteoclast differentiation. Long term follow-up of the critically ill patients after discharge might reveal new insights into the biology of SS and provide fresh directions for therapeutic innovation. Since excessive osteoclast activity contributes to enhanced bone resorption, patients of SS (especially those that survive the acute episode) are likely at risk for bone loss and fracture. Therefore, this pathway is a potential target for therapeutic intervention. There already exists a molecule Simvastatin (*25*), which inhibits osteoclast differentiation by scavenging reactive oxygen species. Zinc is known to inhibit osteoclast differentiation in some other model systems (*12, 27*). It is interesting that prophylactic zinc supplementation improves treatment outcome in human infants aged 7–120 days with probable serious bacterial infection (*4*). We think that zinc supplementation might also play a role in lowering the osteoclastogenic bone loss in SS patients, which might contribute to patient survival and long term recovery (*12*). Antioxidants could also be useful in limiting osteoclast differentiation. However, the value of any therapeutic intervention on the pathway needs to be ascertained through independent testing. Lastly, we have shown in our study that there are 25 genes of the pathway which are significantly differentially expressed in SS (Supplementary Table S4) compared to controls. Since their protein products are detectable in peripheral circulation, it is plausible to suggest these genes, singly or in combination, as plasma biomarkers of SS.

## Conclusion

Firstly, systematic analysis of multiple data sets enabled identification of the core set of genes that are consistently up-regulated in SS. Secondly, we applied three different methods (Over-Representation Analysis, Gene Set Enrichment Analysis and Signaling Pathway Impact Analysis) to arrive at the pathway hsa04380 (Osteoclast Differentiation) with consistent and significant up-regulation in SS. We validated this result with additional bioinformatic analysis and pathway gene expression assay in an independent cohort of SS. Thirdly, we established that the expression signature is distinct in SS compared to control. The 25 genes short-listed, either singly or in combination, may serve as an expression signature of SS. It should be noted that, we do not claim causality, but simply an association of up-regulation of hsa04380 with SS. However, in the light of this finding, altered osteoclast differentiation in septic shock deserves greater attention.

## Competing interests

The authors declare that they have no competing interests.

## Author’s contributions

S.K.M. conceived the study; S.K.M., S.B., S.M. and A.B.P. conducted the data analysis; S.M. conducted the experiment; B.K.D. and P.T. recruited the patients in the validation study; S.K.M, B.R., S.M., B.K.D., P.T. interpreted the results; S.M., S.B. and S.K.M. wrote the manuscript. All authors reviewed the manuscript.

## Acknowledgements

We are grateful to the patients of SCB Medical College Hospital for contributing blood samples for the study. We thank the Director, NIBMG, India for facilitating this work. This study is supported by an intramural grant from NIBMG, Kalyani and an extramural grant (No. BT/PR5548/MED/29/571/2012, duration of 3 years, sanctioned on 27-05-2013) from the Department of Biotechnology, Govt of India. S.M. acknowledges research fellowship provided by the University Grants Commission of India. We thank our colleagues at the NIBMG core facility and the ILS laboratory for timely processing of the samples, often at short notice. S.K.M. acknowledges very helpful advice received from Prof. Gagandeep Kang, Christian Medical College, Vellore during data analysis.

## Supplementary Data

**Supplementary Fig.S1:**
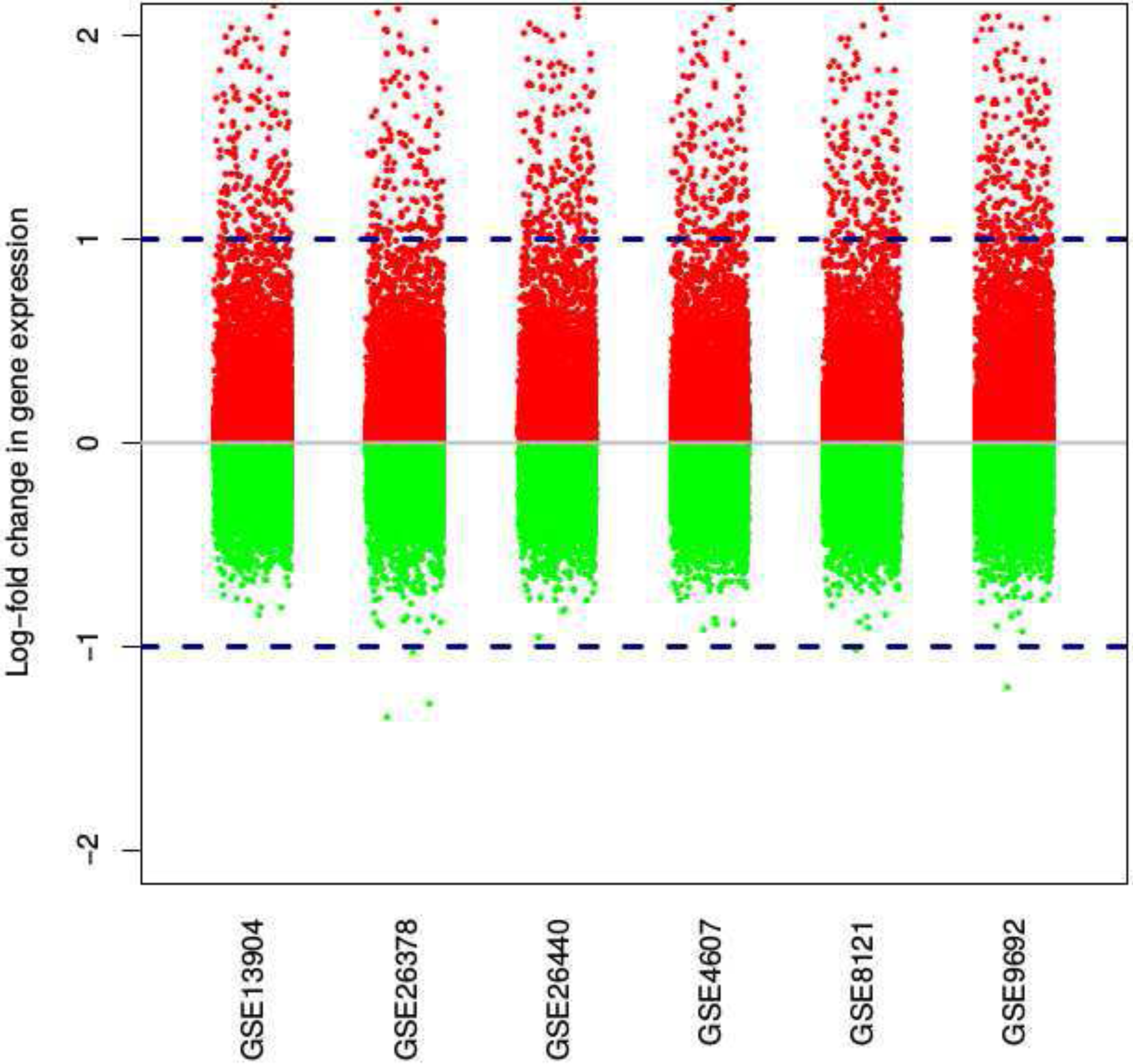
This is a dot plot representing log fold change in gene expression for all the filtered and quantile-normalised genes of each study. Green and Red dots represent down-regulated and up-regulated genes respectively. Any up-regulated gene, i.e., with expression higher in SS compared to healthy controls is shown as a red point, and a down-regulated gene is shown as a green point. The base line at 0 corresponds to “no change” in gene expression. The 2 dotted lines correspond to the threshold of 2-fold change (i.e., log-fold change of 1) in expression. Any point above the top dotted line represents a gene whose expression level is twice or more in SS compared with healthy controls. Similarly, any point below the bottom dotted line represents a gene whose expression level is at least half (or lower) in SS compared with healthy controls. Note that a change of 2 in the linear scale corresponds to a change of 1 in the log-2 scale. There are far more up-regulated genes (red points) than down-regulated genes (green points). Additionally there are many up-regulated genes with 2-fold or greater change in gene expression. This figure provides compelling visual evidence of high-intensity gene up-regulation in SS.

**Supplementary Fig.S2:**
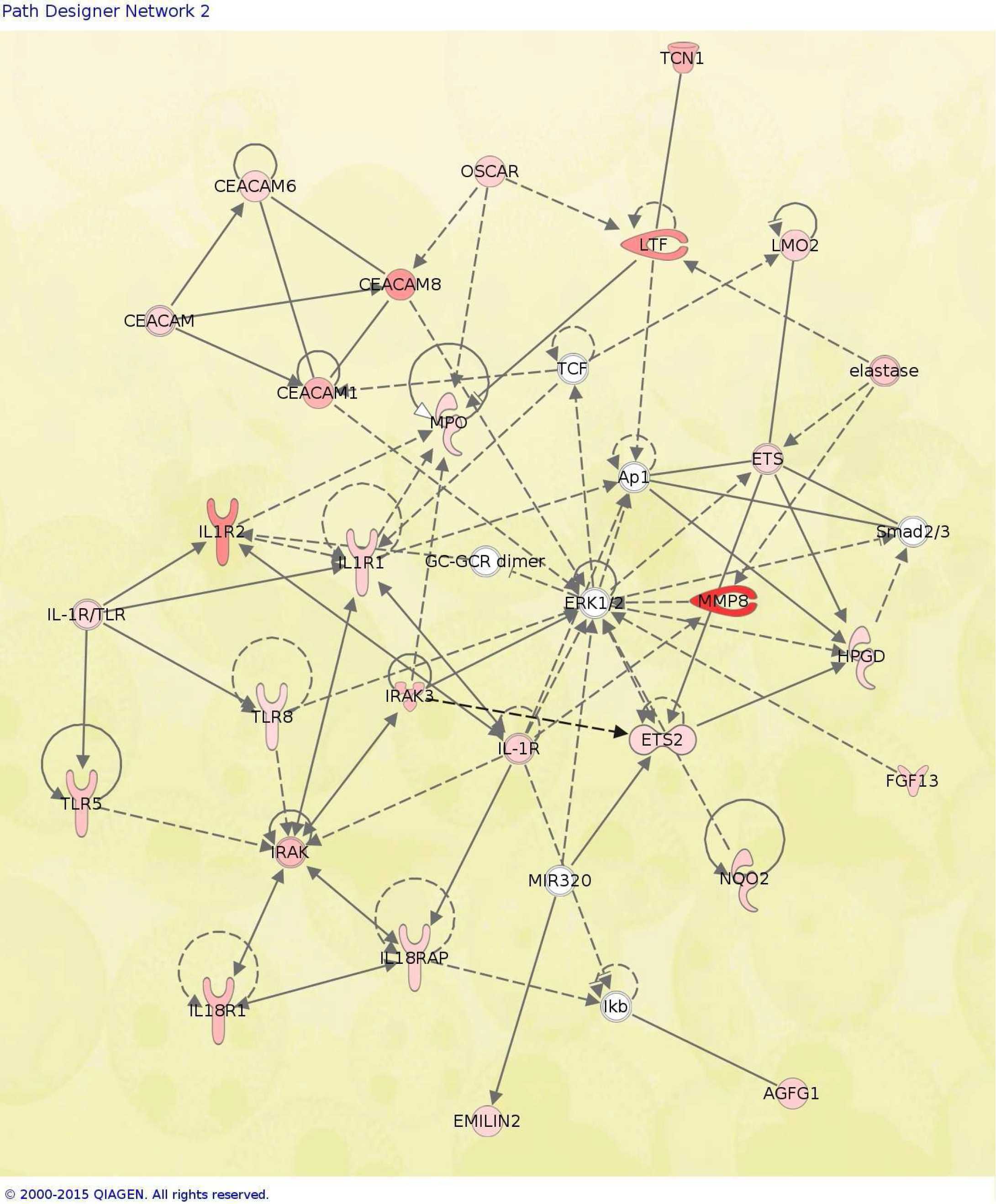
Network “Infectious Disease, Inflammatory Response, Connective Tissue Disorders”, generated by Ingenuity Pathway Analysis with top 200 up-regulated genes with mean log-fold change in gene expression.

**Supplementary Fig.S3:**
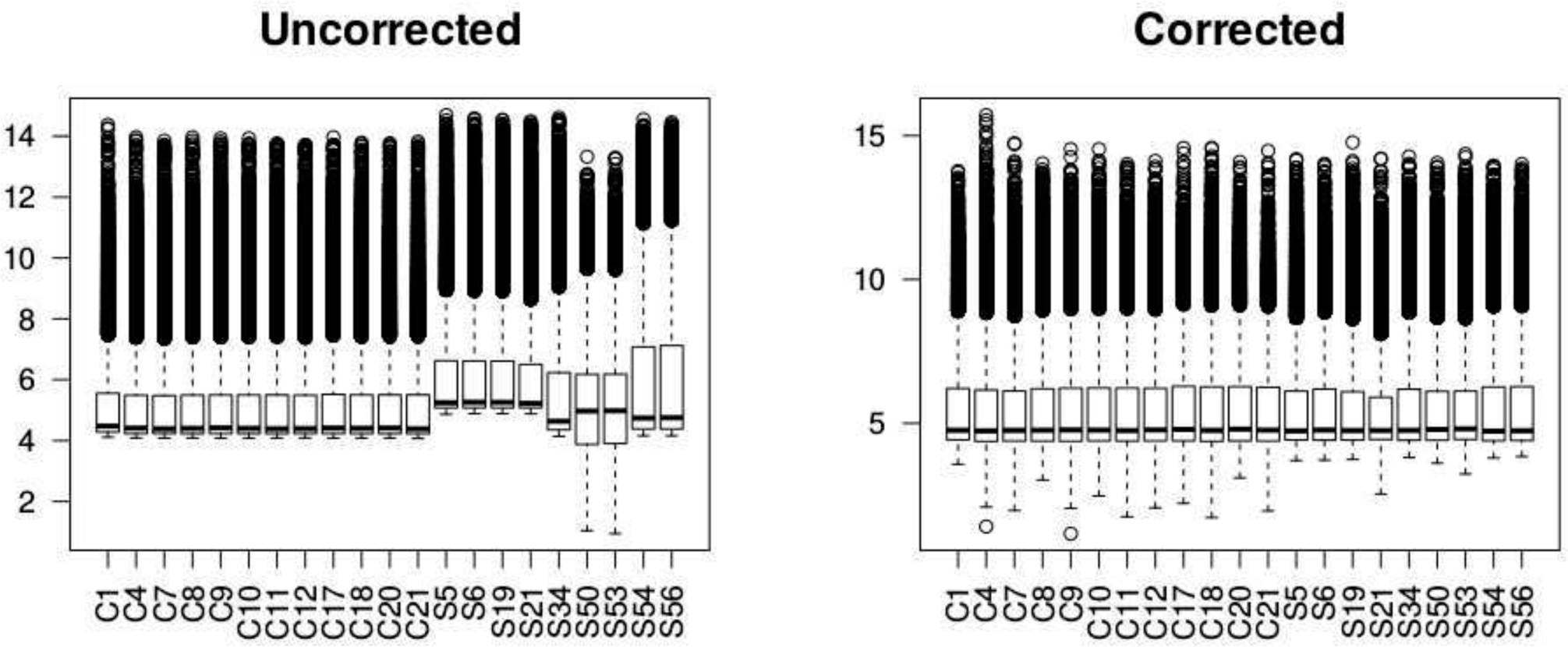
ComBat function (of sva package) was used to remove the batch effect from validation cohort samples as shown in the box plots above, from left(Uncorrected) to right (Corrected).

**Supplementary Fig.S4:**
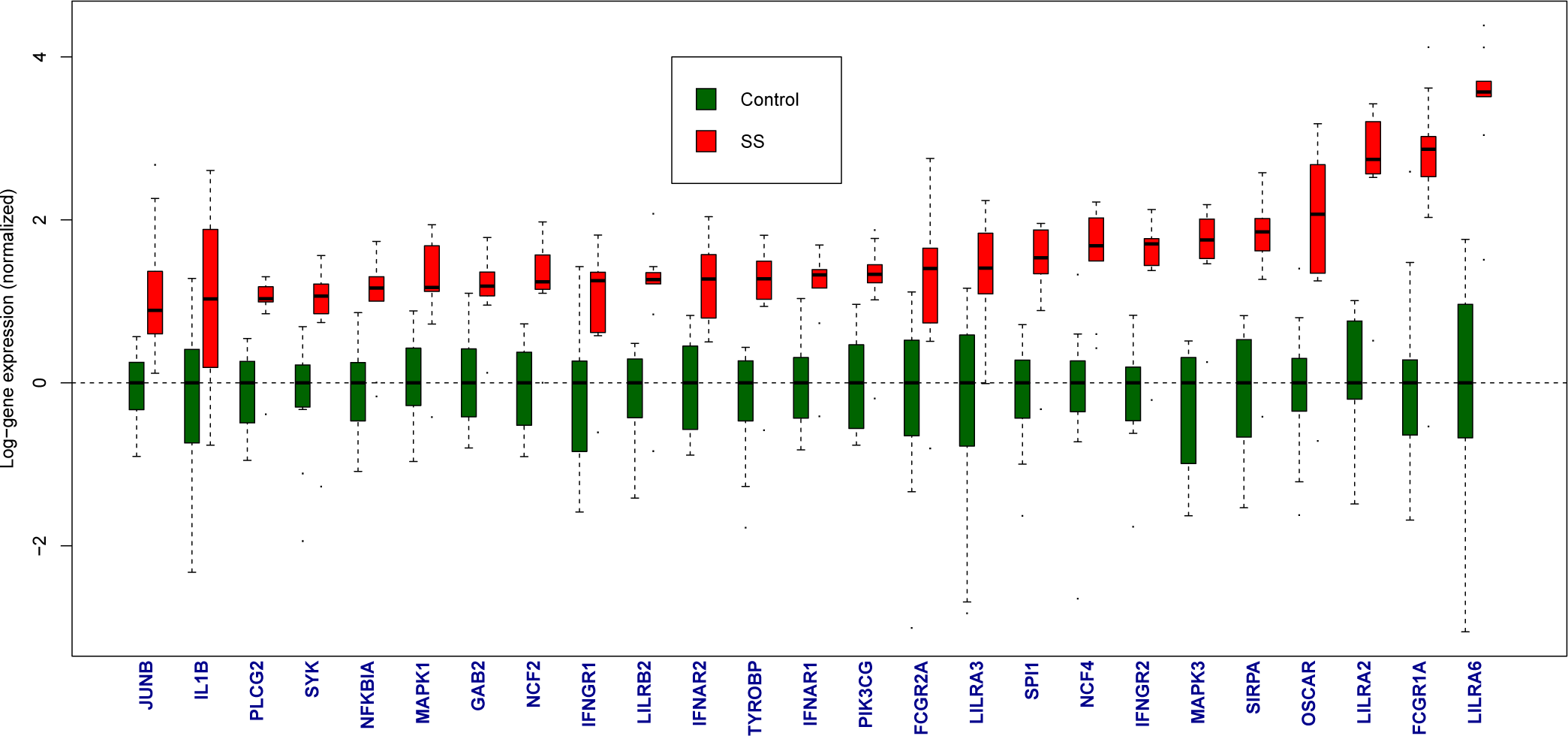
Box plots of the highly significant 25 genes of the pathway hsa04380 up-regulated in SS. Green color corresponds to the control subjects while the red color corresponds to the cases of SS. Gene symbols are shown at the bottom. For each gene, log-intensity of gene expression has been normalized to the median expression of the control group.

**Supplementary Table S1:** Annotation of top 200 genes identified after ORA, as significantly up-regulated in septic shock into associated principal biological functions.

| Biological function | Genes |
| --- | --- |
| Septic Shock<sup>[1],[2]</sup> | ANKRD22, CEACAM1, CLEC5A, G0S2, BCL-6, MMP8, TREM1, CCL4 |
| Inflammation & Innate Immunity<sup>[3]</sup> | SERPINB1, S100A12, SLC11A1, TLR5, PADI4, CLEC5A, TLR8, LILRA5 |
| Anti Microbial Activity<sup>[3]</sup> | ADM, BP1, DEFA4, RNASE3 |
| Complement System<sup>[3]</sup> | C1QB, F5, CR1, C3AR1 |
| Coagulation System<sup>[3]</sup> | CR1, C3AR1, CD59, SERPINA1 |
| Carbohydrate Metabolism<sup>[4]</sup> | PFKB2, PFKB3, SLC2A3, GYG1, PYGL |
| Lipid Metabolism<sup>[5]</sup> | ACSL1, HGF, FAR2, DGAT2 |
| Bone Metabolism<sup>[3]</sup> | CPD, MAPK14, FOSL2, IL1R1, MAP2K6, SOCS3, OSCAR, SIRPA, ALPL, BST1, CA4, TNFAIP6 |
| Calcium Signalling<sup>[3]</sup> | S100A12, DYSF, MRVI1, LILRA5 |
| Cytoskeletal And Cell Adhesion<sup>[3]</sup> | LIMk2, DSC2 |
| Cancer-associated Genes<sup>[3]</sup> | BCL2A1, BCL6, CEACAM1, CD63, DACH1, DSC2, ELANE, ETS2, HPGD, IL1RN, LMO2, MMP8, MMP9, MPO, NQO2, IL1R2, PLXNC1, OLFM4, RAB20 |

---

**Supplementary Table S2:** Top 3 enriched KEGG pathway selected after Over Representation Analysis with top 200 genes.

| KEGG ID | p | odds ratio | Pathway Name |
| --- | --- | --- | --- |
| hsa04610 | 0.00001 | 11.5 | Complement and coagulation cascades |
| hsa04380 | 0.00006 | 6.59 | Osteoclast differentiation |
| hsa05202 | 0.00046 | 4.77 | Transcriptional misregulation in cancer |

**Supplementary Table S3:** Clinical characteristics of SS patients in Validation Cohort.

| ID | Age | SEX | Primary Diagnosis | Vasopressors | Infective focus |
| --- | --- | --- | --- | --- | --- |
| 5 | 70 | M | Septic Shock | Dopamine | NA |
| 6 | 55 | F | Septic Shock | Dopamine | Right middle lobe pneumonia |
| 19 | 14 | F | Septic Shock | NA | Right lower lobe pneumonia |
| 21 | 20 | M | Septic Shock | NA | NA |
| 34 | 32 | M | Septic Shock | Dopamine | Left lower lobe pneumonia |
| 50 | 38 | M | Septic Shock | Noradrenaline | Right leg abscess |
| 53 | 75 | M | Septic shock | Dopamine | Right lower lobe pneumonia |
| 54 | 38 | F | Septic Shock | Noradrenaline | Right lower lobe pneumonia |
| 56 | 63 | F | Septic shock | Dopamine | NA |

**Supplementary Table S4:**
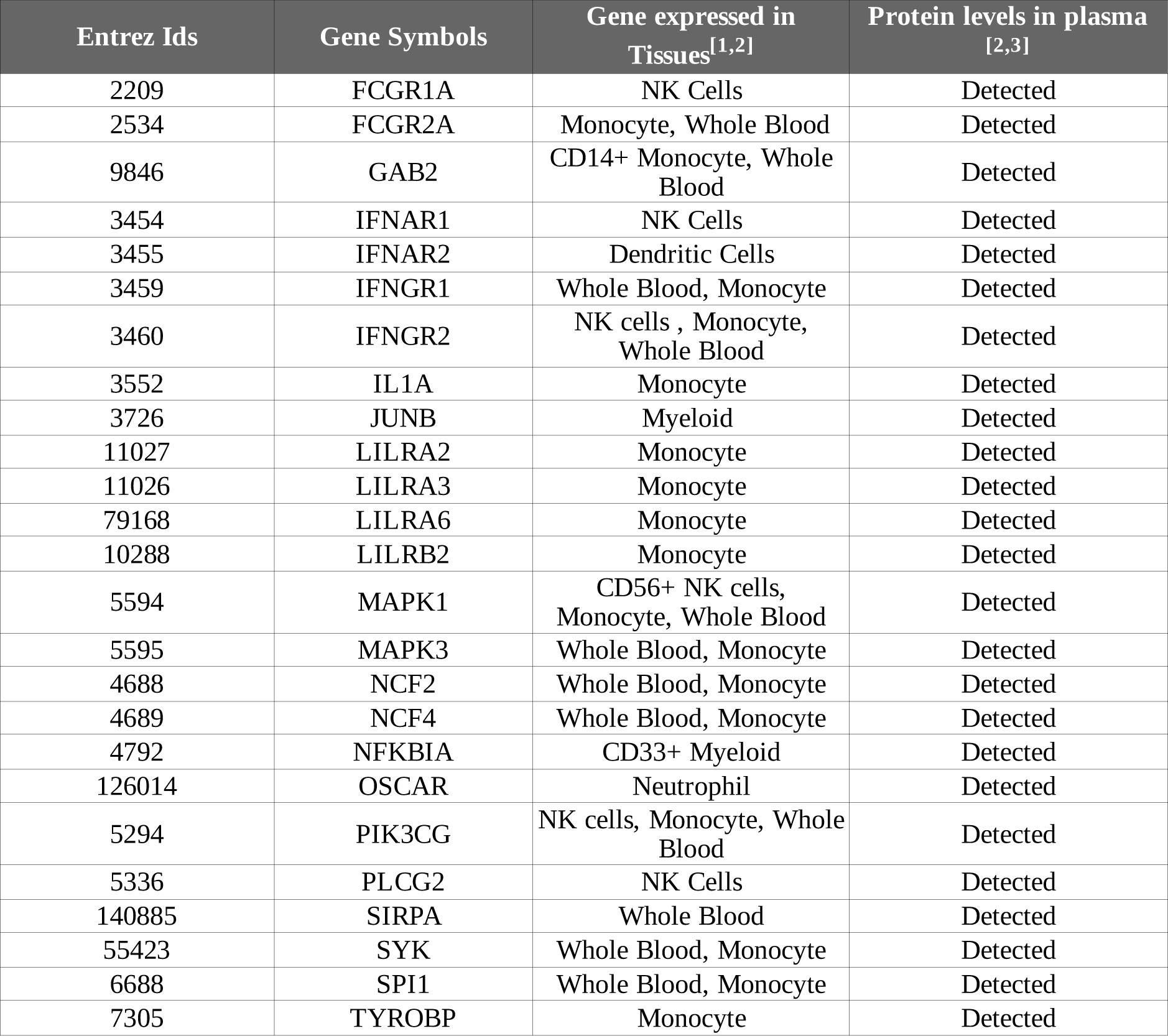
Annotation of 25 genes of the pathway hsa04380 that are significantly up-regulated in SS.

[1] http://www.biogps.org

[2] http://www.genecards.org/

[3] http://www.plasmaproteomedatabase.org/

